# Chemical defences indicate distinct colour patterns with reduced variability in aposematic nudibranchs

**DOI:** 10.1101/2023.01.30.525844

**Authors:** Cedric P. van den Berg, Matteo Santon, John A. Endler, Leon Drummond, Bethany R. Dawson, Carl Santiago, Nathalie Weber, Karen L. Cheney

## Abstract

The selective factors that shape phenotypic diversity in prey communities with aposematic animals are diverse and coincide with similar diversity in the strength of underlying secondary defences. However, quantitative assessments of colour pattern variation and the strength of chemical defences in assemblages of aposematic species are lacking. We quantified colour pattern diversity using Quantitative Colour Pattern Analysis (QCPA) in 13 Dorid nudibranch species (Infraorder: Doridoidei) that varied in the strength of their chemical defences. We accounted for the physiological properties of a potential predator’s visual system (a triggerfish, *Rhinecanthus aculeatus*) and modelled the appearance of nudibranchs from multiple viewing distances (2cm and 10cm). We identified distinct colour pattern properties associated with the presence and strength of chemical defences. Colour patterns were also less variable among species with chemical defences when compared to undefended species. This confirms correlations between secondary defences and diverse, bold colouration while showing that chemical defences coincide with decreased colour pattern variability among species. Our study suggests that complex spatiochromatic properties of colour patterns perceived by potential predators can be used to make inferences on the presence and strength of chemical defences.

## 1. Introduction

Many animals use aposematic colour patterns to warn potential predators of underlying defences [1], with aposematic species in prey communities exhibiting a remarkable diversity of primary (i.e., colour patterns) and secondary defences (i.e., secondary metabolites) [2–4]. However, mechanisms shaping diversity within and among aposematic species in prey communities are complex, and it is poorly understood how the presence and strength of secondary defences correlate with phenotypic diversity in a natural prey community (see [5,6] for discussion). Factors shaping within-species diversity tend to coincide with factors affecting among-species variation in aposematic species (e.g. [7]). This complex mixture of selective mechanisms in natural systems makes it challenging to understand the relationships between primary and secondary defences in prey communities.

Stabilising selection is a crucial driver underlying the distinct appearance of a given aposematic species in a species community. Once aposematism has evolved, stabilising selection is expected to constrain colour pattern diversity within species and Mullerian mimicry rings as predators learn to associate a visual signal with unprofitability [8–13]. Specifically, an invariant appearance across aposematic individuals may facilitate and strengthen predator learning and memorisation. In contrast, variation in signal design may cause predators to make errors when attacking prey and decrease rates of predator learning and increase rates of forgetting [10,14–16]. However, colour pattern diversity within and among aposematic species is ubiquitous. It is thought to be driven by countervailing evolutionary and ecological factors such as genetic drift, gene flow, variation in resource abundance, variation in predator species, and environmental biotic and abiotic variability at different spatial and temporal scales [5,10,17,18]. Aposematism in a spatially homogeneous and temporally stable environment coincides with selection towards reduced colour pattern variability within a population (e.g. [19,20]). In contrast, variability of biotic (e.g. predators) and abiotic factors (e.g. temperature) at spatial and temporal scales can favour selection on phenotypic diversity within aposematic species (e.g. [21–24]) as well as among them (e.g. [25,26]).

Investing in chemical defences is costly (see [6,27] for review) and, as a result, can favour the evolution of various forms of mimicry among prey species (e.g. [28]). Mimicry leads to specific, general or partial (e.g. [29–33]) resemblance among species, reducing phenotypic diversity among chemically defended species and undefended mimics. However, key innovations such as chemical defences are thought to enable niche expansions and, as a result, facilitate speciation [25,34–36]. Adapting to diverse ecological niches, in turn, may lead to phenotypic diversity among aposematic species, especially if such niche specialisations underly changes in the signalling environment, such as the distinctiveness from background habitats or signalling in differing light environments. Indeed, a distinct appearance not only from the background, but also from conspecifics, may aid predator learning [37] and can provide a mechanism to defend against the parasitic effects of certain types of mimicry, such as Batesian and quasi-Batesian mimicry [38–41]. However, long-standing predictions of the benefit of distinctiveness among aposematic species (e.g. [42,43]) are mainly theoretical, with no known studies investigating correlations between distinctiveness and secondary defences among aposematic species in nature.

Attacking well-defended prey is also costly; therefore, predators may generalise more broadly between the colour patterns of previously attacked prey and the prey they subsequently encounter, likely confounded by the cost of making an error (e.g. [44–46]). However, how predator generalisation between and within aposematic species and their mimics influences correlations between secondary defences and colour pattern diversity is complex, highly debated and likely varies among taxa (see [5,6] for discussion). Furthermore, selection for or against colour pattern variability within and among species can act on individual colour pattern elements or perceptual properties rather than the entire animal, depending on which elements of the signal predators learn or pay attention to (e.g. [47]). Therefore, animal colour patterns should be considered complex multicomponent phenotypes [48] under multiple selective pressures (e.g. [48,49]).

When interpreting the ecological relevance of phenotypic variation, it is essential to consider how the appearance of an organism’s colours and patterns change as a function of observer acuity and viewing distance [50]. For example, colour patterns may be cryptic when viewed from a distance but may become aposematic as a predator approaches [50,51]. Animals detect objects and decide their identity and quality based on varying combinations of spatiochromatic features [52–56]. Consequently, predator learning of associations between primary and secondary prey defences, or the subsequent retrieval of formed associations from memory, might happen at a specific range of viewing distances concerning specific spatiochromatic properties of prey appearance. However, the scarce empirical evidence on the ecological significance of colour pattern variability in aposematic animals remains restricted to investigations of colour alone and do not account for the visual acuity of ecologically relevant observers and viewing distance (e.g. [57,58]).

Here, we examined how highly defended aposematic nudibranch species differ from less well-defended species in appearance to a potential predator and if, among species, variation in perceived colour patterning varies with the presence and strength of chemical defences. Specifically, we hypothesised that chemical defences would correlate with increases or decreases in colour pattern distinctiveness between species as perceived by a potential predator. We further hypothesised that colour patterns in chemically defended species were less variable than in species without chemical defences as perceived by a potential predator. To do this, we modelled the visual appearance of 13 sympatric Dorid nudibranch species across multiple viewing distances corresponding to the later stages of an escalating predation sequence [14,59]. We quantified the perception of within-species colour pattern variability using the Quantitative Colour Pattern Analysis (QCPA) [60], allowing for the consideration of colour, luminance and spatial vision of triggerfish (*Rhinecanthus aculeatus*). Using exploratory factor analysis, we identified latent variables to compare the colour pattern appearance of individuals belonging to three levels of chemical defence. Chemical defences were defined using previously published assay data [61,62]. We then investigated differences in the perceived appearance and variability of colour patterns for species belonging to each level of chemical defences.

## 2. Materials and Methods

### (a) Study species

We used digital photographs of 311 Dorid nudibranchs using a calibrated Olympus EPL-5 with a 60mm macro lens (see the Supplement for details on camera calibration). These individuals belonged to 13 species: *Aphelodoris varia* (N=31), *Chromodoris elisabethina* (N=31), *Chromodoris kuiteri* (N=49), *Chromodoris lochi* (N=8), *Chromodoris magnifica* (N=14); *Dendrodoris krusensterni* (N=7); *Discodoris* sp. (N=10); *Doriprismatica atromarginata* (N=35); *Glossodoris vespa* (N=32); *Hypselodoris bennetti* (N=13); *Phyllidia ocellata* (N=32)*, Phyllidia varicosa* (N= 9), *Phyllidiella pustulosa* (N=40) (Fig. 1) from five locations on the east coast of Australia: Mackay (QLD), Sunshine Coast (SE Queensland, QLD), Gold Coast (SE QLD), Cook Island (New South Wales, NSW) and Nelson Bay (NSW) between March 2016 and February 2021. Two out of 13 species (*Doriprismatica atromarginata, Goniobranchus splendidus*) were sampled across sites in QLD and NSW in high numbers, whereas the other species were only sampled in either NSW or QLD or with highly uneven numbers between sites (Table S1). Two individuals of *Chromodoris magnifica* were provided by an aquarium supplier (Cairns Marine, Pty Ltd, Cairns, QLD). These species were selected as they were relatively abundant at our study sites and covered a broad range of visual appearances and strengths of chemical defences. Furthermore, we have previously provided data on the strength and identity of chemical defences in these species sampled from the same locations as individuals from this study [61,62].

**Figure 1.**
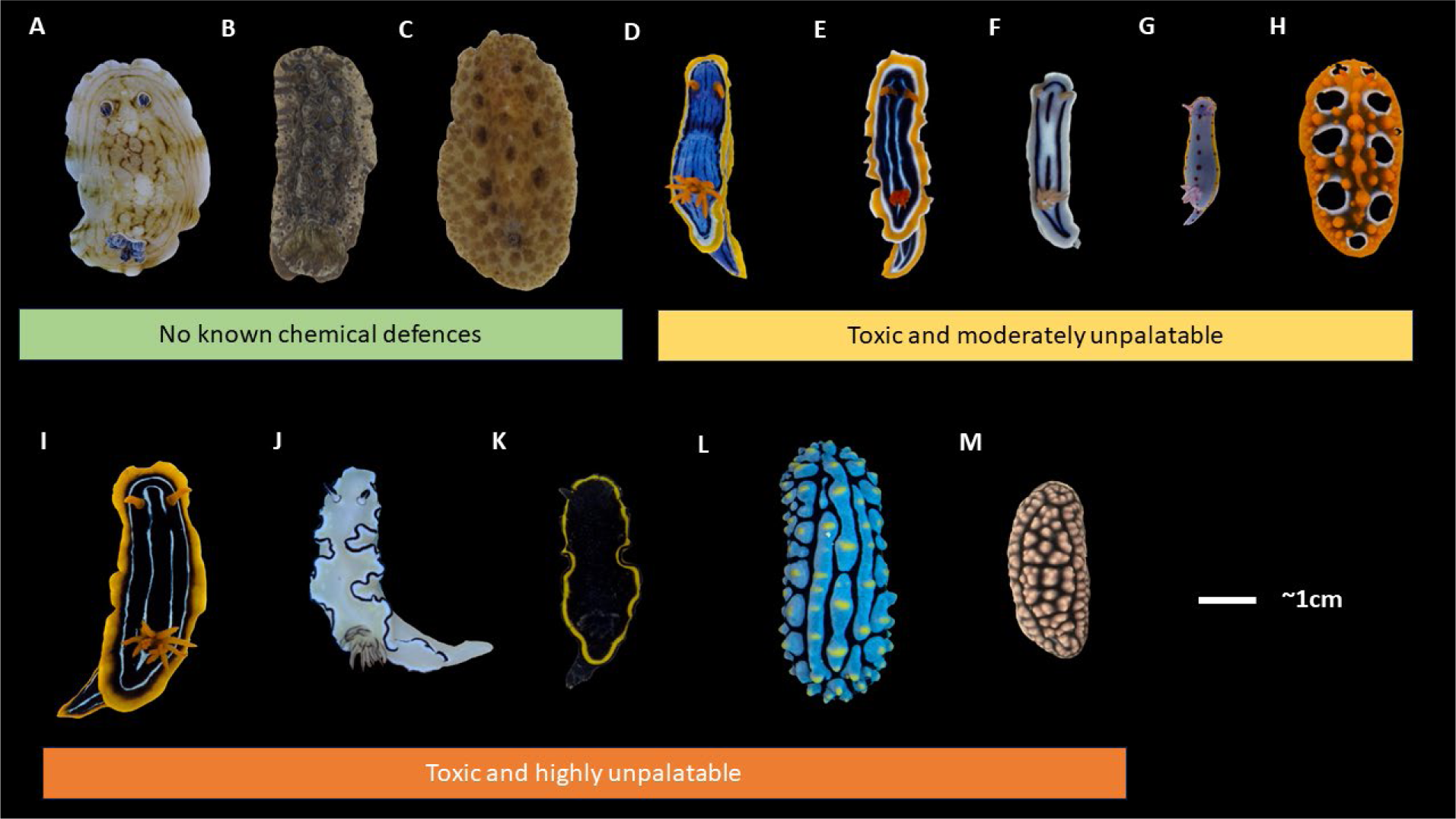
Representative photographs of the 13 species used in this study grouped into categories of chemical defences based on whole-body extract assays with palaemon shrimp to assess unpalatability (1-Effective Dose, ED_50_) and brine shrimp to assess toxicity (1-Lethal Dose, LD_50_) values as per [61,62]: A) *Aphelodoris varia;* B) *Dendrodoris krusensterni*; C) *Discodoris sp*; D) *Chromodoris elisabethina*; E) *Chromodoris magnifica;* F) *Chromodoris lochi;* G) *Hypselodoris bennetti;* H) *Phyllidia ocellata*; I) *Chromodoris kuiteri*; J) *Doriprismatica atromarginata;* K) *Glossodoris vespa*; L) *Phyllidia varicosa;* M) *Phyllidiella pustulosa*.

Most nudibranchs were photographed underwater against their natural habitat (n = 182) with the camera in an Olympus PT-EP10 underwater housing and using white LED illumination from a combination of VK6r and PV62 Scubalamp video lights. The remaining nudibranchs (n = 129) were collected for separate studies on their chemical defences, taken back to the laboratory, submerged in water in a petri dish and photographed against a white background with the same camera. In the laboratory, nudibranchs were illuminated with 400nm-700nm full-spectrum white LED lights. The Supplementary Information (Table S1) details collection sites and dates, and camera and illumination spectra are provided in [60]. A sub-sample of these images was previously used to investigate distance-dependent signalling regarding colour pattern detectability and boldness [63]. Nudibranchs were collected under the Queensland General Fisheries Permit 183990, 205961 and NSW Scientific Collection Permit P16/0052-1.0.

### (b) Image analysis

We used ImageJ [64] and the MICA toolbox [65] to manually segment the images into regions of interest (ROI). This was done by outlining and selecting the animal from its background and defining a size standard. All nudibranchs were aligned head up in the image before analysis with QCPA [60], with the rotation angle determined by the rotation, causing most of each animal to be aligned vertically. To analyse the nudibranch colour patterns, we used the visual system parameters of a trichromatic triggerfish, *Rhinecanthus aculatus* [66–71], a common shallow reef inhabitant found throughout the Indo-Pacific, which feeds on invertebrates, algae, and detritus [72].

We analysed colour patterns for viewing distances of 2cm and 10cm, using the estimated spatial acuity of the triggerfish of three cycles per degree [66,70]. A viewing distance of 2cm represents the spatiochromatic information available to a triggerfish upon immediate contact with a nudibranch. A viewing distance of 10cm more likely represents visual information available to a triggerfish at a short distance where a subjugation attempt has not yet been made. Following acuity modelling, the images were processed with a Receptor Noise Limited (RNL) ranked filter (falloff: 3, radius: 5, repetition: 5) and clustered using RNL clustering with a colour threshold of 2 *ΔS* [71,73] and a luminance contrast threshold of 4 *ΔS* [74] for all analyses except the local edge intensity analysis (LEIA) which does not require RNL clustering but is recommended to be subjected to RNL ranked filtering [60]. We calculated receptor-specific Weber fractions based on a relative photoreceptor abundance of 1:2:2:2 (sw:mw:lw:dbl) and photoreceptor noise of 0.05, resulting in 0.07:0.05:0.05:0.05.

QCPA analysis was achieved using a custom batch script [75] running on high-performance computing (HPC) infrastructure. We analysed each animal colour pattern using:

1. colour adjacency analysis (CAA), which describes pattern geometry in a segmented image;
2. visual contrast analysis (VCA), which describes pattern boldness based on chromatic and spatial pattern element properties in a clustered image; 3) boundary strength analysis (BSA), which describes the colour and luminance contrast of boundaries between pattern elements at the scale of an animal in an unclustered image; and 4) local edge intensity analysis (LEIA) which describes the strength of colour and luminance contrast at the scale of an edge-detecting receptive field in an unclustered image. This resulted in a highly descriptive array of 157 colour pattern statistics for each animal. A detailed description of each pattern statistic can be found in [60]. Here, we use CAA, VCA, BSA, and LEIA as prefixes for each type of analysis.

All pattern analyses, except LEIA, used a segmented image and measured transitions between pixels along vertical (along body axis) and horizontal (perpendicular to body axis) sampling transects in a transition matrix. Statistics ending with ‘vrt’ or ‘hrz’ are the vertical (i.e., up-down in image) and horizontal version (analysing the respective transition matrix only) of their respective statistic (analysing the full transition matrix) and can represent differential directionality sensitivity in the visual system of an observer and directionality in patterns such as stripes [76–78]. LEIA does not use a transition matrix due to the lack of image segmentation but equally discriminates between horizontal and vertical edge contrast by describing the shape of a histogram drawn from edge contrast measurements in a given image or region of interest [60].

### (c) Chemical defences

To categorise the level of chemical defences for each species, we used previously published data on the deterrent properties from feeding rejection assays with rockpool shrimp (*Palaemon serenus*), which demonstrate similar results to assays performed with triggerfish and toadfish [61] and toxicity assays with brine shrimp [61,62]. Assays were conducted by adding extracted nudibranch compounds to food pellets made from squid mantle at increasing concentrations. Effective dose (ED_50_) and lethal dose (LD_50_) values in [61,62] were calculated based on the concentration that elicited a rejection response in, or mortality of, at least 50% of the shrimp. For this study, we averaged ED_50_ and LD_50_ values from [61] when multiple extracts from the same species were reported. We considered only whole-body extracts (rather than mantle-only values) to make assay values comparable between species. We then subtracted these values from 1 so that values close to 0 were the most palatable/non-toxic, and values close to 1 were the least palatable/ toxic (Table S2). Although *C. magnifica* was not included in [61], [79] demonstrated that this species also stores latrunculin A as the sole defensive compound in the mantle rim, and this is at concentrations between those found in *C. elisabethina* and *C kuiteri* [80]. We, therefore, set unpalatable ED_50_ values as the average from these two sister species for *C. magnifica*. Lastly, assay data for *G. vespa* is presented in [62].

Like Winters et al. [61], we binned the species into categories indicating chemical defence strength to account for our dataset’s highly uneven spread in toxicity and palatability values and the difference in sampling levels between colour pattern data and chemistry data. Our categorisation differed from that of Winters et al. [61] in that we based our categories on the assumption of a sigmoidal dose-effect response similar to a psychometric curve. Species were allocated in the following classes (Fig. 1), where we treated NR values from [61] as 0:

1.) Not defended (1 - ED_50_ / LD_50_ = 0)

2.) Toxic and moderately unpalatable (0.25 < 1 - ED_50_ > 0.74 and LD_50_ > 0),

3.) Toxic and highly unpalatable (0.74 < 1 - ED_50_ and LD_50_ > 0).

The threshold to distinguish between medium and high levels of unpalatability was 0.74, representing the median 1 – ED_50_ value of chemically defended species while also being very close to the point-of-inflexion in a sigmoidal response curve. Only 3 out of 10 species with chemical defences had 1 – LD_50_ values below 0.5, yet 6 out of 10 had values above 0.80. Therefore, we did not distinguish between different toxicity levels in our dataset. Treating toxicity as present/absent and distinguishing between medium and high levels of unpalatability ensured at least three species in each category, allowing the investigation of differences in animal colouration between variable levels of chemical defences.

### (d) Statistical analysis

Our study considers many of the more commonly found Dorid nudibranchs in the study sites (e.g. [22–24]). To analyse the large dataset derived from the QCPA analysis, we only kept images that did not produce any missing value for any pattern metrics. VCA, CAA, and BSA metrics can produce NaN or infinite values if a colour pattern has less than two colour pattern elements following RNL clustering [60]. LEIA metrics do not suffer from this limitation. Nine available images from *Discodoris sp* were rejected from analysis due to this constraint, resulting in the reported sample size.

We then applied a Latent variable Exploratory Factor Analysis (EFA) with the R package *psych* using the factoring method of Ordinary Least Squares ‘ols’, and the orthogonal rotation ‘varimax’. To prepare the dataset for the EFA, we first filtered the number of highly correlated QCPA metrics by keeping only those that were less correlated than 0.6 (Pearson correlation), which reduced their number from 157 to 15. We then run the factor analysis based on three factors. The number of factors was selected by comparing the eigenvalues calculated from the original dataset to the median eigenvalues extracted from 10,000 randomly generated datasets with the same number of rows and columns of the original data. We selected factors with eigenvalues greater than the median of the eigenvalues from the simulated data. We also computed bootstrapped confidence intervals of the loadings by iterating the factor analysis 1000 times.

Looking at the loadings of each factor, we can identify what latent variable they describe. While it would be possible to discuss each factor extensively, we keep their description to loadings of +/- 0.4 (out of 0 -1) to capture their main properties. Due to data filtering for metrics less correlated than 0.6, the QCPA parameter listed for a given loading is likely synonymous with various other parameters in our 157-dimension colour pattern space (Table S5). Therefore, the precise wording to describe each factor can vary depending on which colour pattern metrics are put into focus—for example, *BSA.BMSL.Vrt* is positively associated with factor 1 (Fig. 2) but is simply a placeholder for *BSA.BMSL* (both considering horizontal and cumulative transitions) as it is 92-96% correlated with these metrics and 97% correlated with *BSA.BML* (Table S2). Unlike *BSA.BMSL* (which describes boundary contrast using the mean RNL luminance contrast between colour pattern elements relative to the fraction of the respective pattern border), *VCA.BML* captures boundary contrast calculated by the Weber contrast of cone catches in the luminance channel between colour pattern elements relative to the fraction of a given boundary type. Thus, it would be more appropriate to say that animals with high values of factor 1 are associated with stronger achromatic colour pattern boundary contrast rather than explicitly referring to the randomly retained value only. A complete list of all colour pattern parameters with more than 0.6 Pearson correlation with parameters associated with factors 1-3 shown in Fig. 2 can be found in the Supplement (Table S2).

**Figure 2:**
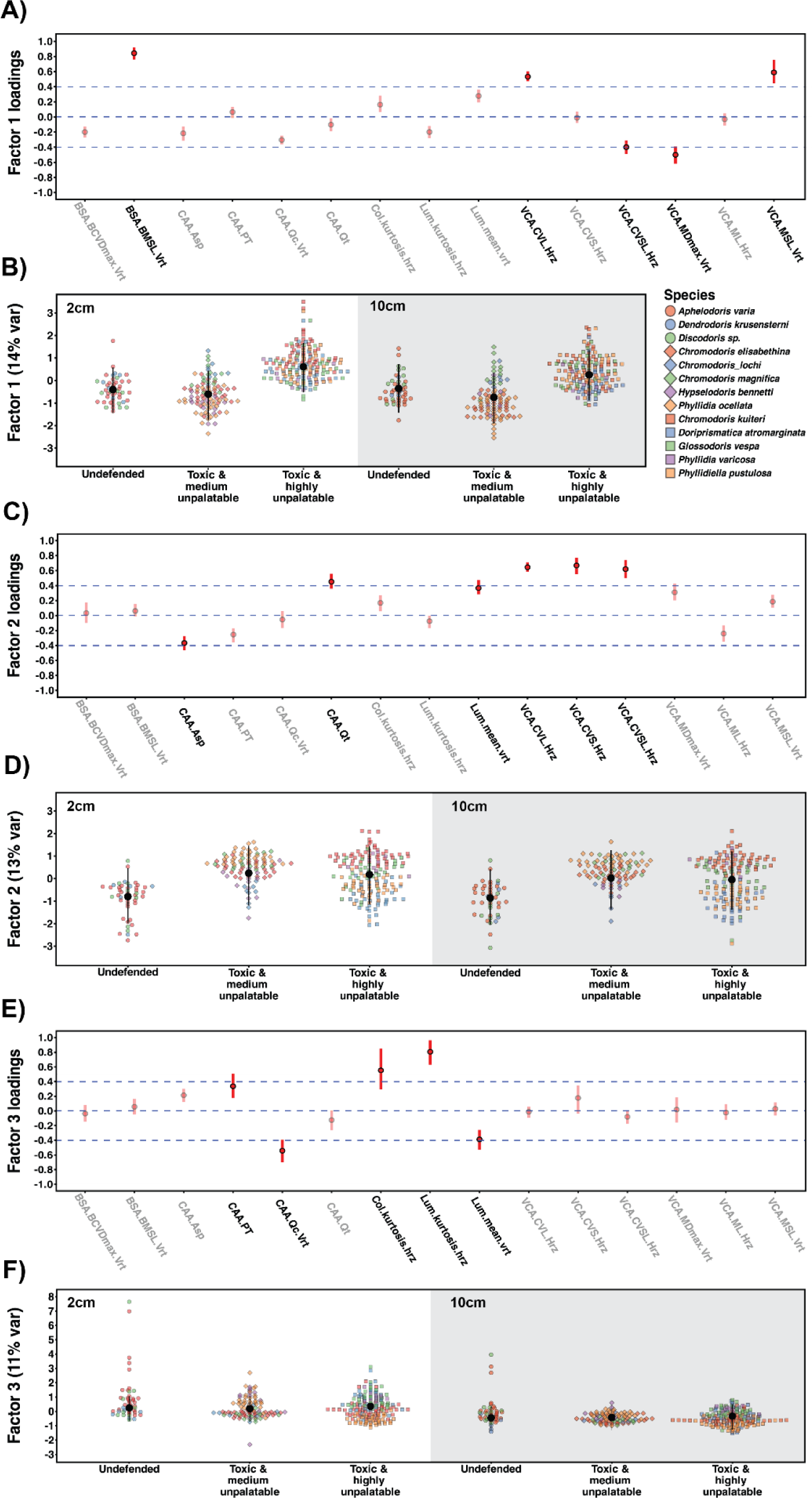
Detailed visual representation of the loadings of factor 1 (A), factor 2 (C) and factor 3 (E). Greyed-out factor loadings indicate colour pattern descriptors with minor contributions to each factor. Factor values for each group with different strength of chemical defences are given for factor 1 (B), factor 2 (D) and factor 3 (F). Estimates are given for 2cm viewing distance (left panel half, white) and 10cm (right panel half, grey). Coloured points represent repeated observations for each species (N = 13). Black vertical bars represent group predicted medians and 95% compatibility intervals (CIs) derived from the joint posterior distributions of the model

The scores of the factors extracted from the EFA were then used to implement three phylogenetic, distributional linear mixed models to compare the colour patterns of nudibranchs with different levels of chemical defences. Models were run in R v 4.1.2 (https://www.r-project.org/) using the brms package [81], which fits Bayesian models using Stan (https://mc-stan.org/). To account for the phylogenetic dependency of closely related species, all models included the phylogenetic tree of the 13 species of nudibranchs (Fig. S1), with the tree from [82] pruned and missing species added according to their taxonomic classification in the World Register of Marine Species [83]. The phylogenetic model was implemented following the guidelines of the online *brms* vignette (https://cran.rproject.org/web/packages/brms/vignettes/brms_phylogenetics.html) based on de Villemeruil & Nakagawa [84].

The first model investigated differences in scores for latent *factor 1* between nudibranchs with different levels of chemical defences (see chemical defences section) using a Student distribution. The model estimated the effect of the main categorial predictors level of *chemical defence* (undefended; toxic and moderately unpalatable; toxic and highly unpalatable) and *observer distance* (2 cm and 10 cm) and their interaction on both the mean and the residual standard deviation of the response distribution. To account for repeated measurements of each species, we also included *species ID* as a random intercept to the model. We further included random slopes over distance because their relationship with the value of the response *factor 1* varied among species. The second and third models were identical to the first but used *factor 2* and *factor 3* as response variables.

All models were fitted using weakly informative prior distributions (normal with mean = 0 and sd = 5 for intercept and coefficients, exponential (1) for standard deviations). Their performance was evaluated using posterior predictive model checking, which compares model predictions with observed data to assess overall model fit. We ran four Markov-Chain-Monte-Carlo (MCMC) chains for each model and obtained coefficient estimates from 16,000 post-warm-up samples. All model parameters reached reliable conversion indicators [85]: A Monte Carlo standard error smaller than 5% of the posterior standard deviation, an effective posterior sample size greater than 10% of the total sample size, and a *R̅* statistic value smaller than 1.01.

We present the medians of latent factors values and their 95% credible intervals of the posterior distributions of fitted values for the population average, obtained from the joint posterior distributions of the model parameters for the combination of chemical defences and distance [85,86] (Fig. 2). The same posterior distribution of fitted values was used to compute pairwise differences and their 95% credible intervals between all the combinations of the same two categorical predictors using the ‘emmeans’ R package [87]. To compare variances of responses between all predictor groups, we also computed the posterior distribution of all pairwise differences of the residual standard deviation on the original scale (back-transformed from the log scale). The effect size of pairwise differences increases with increasing deviation of such differences from zero, and the robustness of the result increases with decreasing degree of overlap of the 95% Credible Intervals (CIs) with zero.

## 3. Results

We identified three latent factors describing overall differences in colour pattern appearance to a triggerfish (*R. aculeatus*). We describe each factor at 2cm and 10cm, respectively.

While not intended to identify a maximal amount of variability in colour pattern variation in our dataset, the three factors still explain 38% of the total variation (factor 1: 14%; factor 2: 13%; factor 3: 11%) (Fig 2).

### (a) <u>Factor 1:</u> Colour patterns with high achromatic contrast have low colour contrast

Contrasts [difference (+-95% CI)] between groups of chemical defences indicate that toxic species with high levels of unpalatability differed in appearance from toxic species with moderate levels of unpalatability (Fig. 2b, Table S3). However, undefended species did not differ from chemically defended species for factor 1. At a 2cm viewing distance, undefended species are not different in appearance from toxic and highly unpalatable species (0.99 (−2.31 / 0.31)). In contrast, toxic and moderately defended species have a lower score (−1.23 (−1.74 / - 0.70)) for factor 1 compared to highly unpalatable toxic species (Fig. 2b). This is true at immediate contact between the triggerfish and prey at 2cm, as well as at 10cm (undefended vs. toxic and highly unpalatable: -0.60 (−2.00 / 0.81); toxic and medium unpalatable vs. toxic and highly unpalatable: -1.01 (−1.67 / -0.33)). Toxic animals with medium levels of unpalatability did not differ from undefended species regarding factor 1 at either 2cm (0.21 (−1.10 / 1.56) or 10cm (0.40 (−1.09 / 1.82). We found no indication of differences in colour pattern variability in species of different groups as captured by factor 1 (Table S4).

Factor 1 describes 14% of colour pattern variability in our dataset. It is associated with high loadings of luminance contrast between colour patches as a function of their patch size, which VCA describes. We can see high loadings for mean and standard deviation variation measures of pattern contrast measured as cone catches of the luminance channel (e.g. *VCA.CVL*) and using the RNL model (e.g. *VCA.MSL*). We also find high luminance pattern contrast captured by factor 1 as an expression of the boundary contrast (BSA), which refers to contrast scaled by the length of boundaries between colour patches rather than their size. Given that larger patches tend to have longer boundaries, it is not surprising that we find similar loadings for measures relative to either. The negative loadings for chromatic colour pattern contrast (e.g. *VCA.MDmax*) indicate that patterns with strong and variable achromatic contrast tend to have a reduced level of average chromaticity contrast. High factor values would indicate the presence of black and white, pale hues or long wavelength colours that appear of low chromaticity to the visual system of a triggerfish. Therefore, our results indicate higher levels of achromatic contrast and lower levels of chromatic contrast present in the colour patterns of highly unpalatable toxic species compared to the other groups, with the increase in achromatic contrast coinciding with more prominent relatively achromatic colour pattern elements.

### (b) <u>Factor 2:</u> Highly contrasting colour patterns are more regular and vertically elongated

Contrasts [difference (+-95% CI)] between the different groups of chemical defences indicate that chemically defended species do not have higher scores for factor 2 than undefended species (Fig. 2d, Table S5). There was also no difference in factor values between toxic and medium unpalatable animals and toxic and highly unpalatable animals at either 2cm (0.05 (−0.84 / 0.94) or 10cm (0.06 (−0.91 / 0.92)). However, at 2cm viewing distance, undefended species have more variable colour patterns than toxic and moderately unpalatable species (0.40 (0.14 / 0.74) as well as toxic and highly unpalatable species (0.31 (0.06 / 0.67)) (Table S6).

Factor 2 explains 13% of colour pattern variability in our dataset. It describes the relationship between decreases in the aspect ratio of colour patterns (*CAA. Asp*) coinciding with decreases in average patch size (*CAA.Pt*) as well as decreases in the average luminance contrast (e.g. *VCA.ML*) and its variability (e.g., *VCA.sL*) between patches in the horizontal axis and increases in various measures of chromatic and achromatic colour pattern contrast variability relative to the mean contrast in a given colour pattern (e.g. *VCA.CVSL, VCA.CVS*) as well as increases in colour pattern transition regularity (e.g., *CAA.Qt*).

### (c) <u>Factor 3:</u> Colour patterns with variable edge contrast have reduced spatial evenness

Contrasts [estimate (+-95% CI)] calculated between the different groups of chemical defences indicate no overall differences between groups (Fig. 2, Table S7). This is the case for both 2cm (undefended vs. toxic and medium unpalatable: 0.05 (−1.16 / 1.14); undefended vs. toxic and highly unpalatable: -0.10 (−1.29 / 1.03); toxic and medium unpalatable vs. toxic and highly unpalatable: -0.16 (−0.77 / 0.46)). We found no indication of differences in colour pattern variability in species of different groups captured by factor 3 (Table S8).

Factor 3 explains 11% of colour pattern variability in our dataset. It describes positive changes in colour (e.g. *Col.kurtosis*) and luminance (e.g. *Lum.kurtosis*) contrast variability relative to the average contrast in an animal coinciding with reduced colour pattern evenness (e.g. *CAA.Q_C_*) as well as decreased average luminance contrast of boundaries between colour pattern elements (e.g. *Lum.mean*) and decreased overall colour pattern complexity (*CAA.C*).

## 4. Discussion

We identified three latent variables that captured differences in appearance between distinct differences in colour patterns between our three levels of chemically defended groups of nudibranch molluscs (Fig. 2). Our analysis captures a significant proportion of variability in the dataset (38%) and indicates substantial colour pattern variation among sampled species across multiple viewing distances as perceived by a potential predator (Fig. 2). We found differences in appearance both between chemically defended and undefended species and also between toxic/moderately unpalatable species and toxic/highly unpalatable species. These differences in colour patterns between species belonging to different levels of chemical defences are likely visible to a potential predator at close contact (2cm) and from further away (10cm) and might be used by predators to infer the presence and strength of underlying chemical defences based on the general appearance of prey animals.

The colour patterns of chemically defended species were less variable than those of undefended species (Fig. 2d, Table S3). Specifically, the variability of colour and luminance contrast and the spatial arrangement of colour pattern elements was reduced in species with chemical defences compared to those without. Furthermore, the colour patterns of toxic species with high levels of unpalatability were different in appearance from toxic species with moderate levels of unpalatability (Fig 2b, Table S3). Specifically, species with high levels of unpalatability showed increased levels of achromatic contrast between colour pattern elements when compared to more palatable toxic species. This increase in achromatic contrast in highly unpalatable species coincides with a decrease in the mean level of chromatic contrast relative to toxic species with lower levels of unpalatability. Overall, the differences in the visual appearance to a potential predator between species of nudibranchs with different levels of chemical defences describe general colour pattern properties (such as pattern regularity and spectral contrast) associated with aposematic signalling (Fig. 2). Therefore, in agreement with existing literature (e.g. [2,88]), we find that Dorid nudibranch colour patterns are highly diverse and that the presence of chemical defences correlates with the presence of boldly contrasting colour patterns.

The observed differences in animal colouration between groups of species with varying levels of chemical defences generally agree with and can be interpreted as indicating selective factors driving between-species pattern diversity in conjunction with the presence of secondary defences. Such drivers of phenotypic diversity can favour distinctiveness among chemically defended species, either as a means to defend against Batesian mimicry (e.g. [38]), as well as the potential need to optimise signalling efficacy across a complex, spatially and temporally variable biotic and abiotic environment (e.g. [5,17,21,22,25,89,90]). Thus, our results agree with predictions made by assuming facilitated niche expansion and subsequent speciation and adaptation to visually diverse habitats [25,34–36] as potential drivers of phenotypic diversity in chemically defended species.

Our results further suggest the general presence of secondary defences to coincide with reduced colour pattern variability among species when viewed up close by a potential predator (Fig. 2e, Table S3). Reduced variability among chemically defended species may suggest the presence of broadly generalisable, qualitative signalling properties underlying aposematic signalling in the species considered in this study. However, the presence of distinct colour pattern appearance at a quantitative scale (i.e., comparing species with different levels of chemical defences) would align with chemical defences, favouring visual distinctiveness from co-occurring Batesian or quasi-Batesian mimics (e.g. [38]). In other words, considering colour patterns as complex, multicomponent signals, it is possible to think of certain colour pattern properties indicating the qualitative presence of secondary defences (‘is the animal defended or not’). In contrast, others indicate the quantitative presence of secondary defences (‘how potent are the defences’), thus allowing different parts of simultaneously perceived visual information elicited by animal colouration to be under seemingly opposing selection pressures towards and away from general resemblance. In addition to these perceptual modalities being realised simultaneously, trade-offs between selective pressures for and against multiple, seemingly contractionary signalling properties of colour patterns can be mediated by distance-dependent signalling (e.g. [63,94]). Our results suggest both to be possible, with colour pattern variability only differing between species with and without chemical defences at 2cm viewing distance but not 10cm. In contrast, toxic and highly unpalatable species differ in their appearance from toxic and moderately defended species as well as undefended ones at 2cm and 10cm.

Phenotypic diversity within (e.g. polymorphism and polyphenism) and among chemically defended species is generally described as a detriment to predator learning, with selection towards resemblance underlying purifying selection at the species level (e.g. [10,14–16]) and Mullerian mimicry at the community level (e.g. [7,32,91,92]). However, phenotypic diversity among chemically defended species might, contrary to general assumptions, benefit predator learning as it can lead to more stable, generalisable associations [93] and, thus, provide mutual benefits among chemically defended species considered in the context of qualitative and quantitative signal honesty and mimicry. Experimental investigations into the importance of signal variability for avoidance learning in non-human animals would be of great interest for future research as it, in turn, would inform our assumptions on the mechanisms underlying the evolution and maintenance of colour pattern diversity within and among chemically defended species.

Our methodology is tailored to reflect the fact that colour pattern elements and signalling properties do not exist in isolation, thus warranting an ‘agnostic’ approach to deduce correlations between predictor and dependent variables in the context of a complex trait described by a high-dimensional dataset (i.e., colour pattern space) [55,60,95]. Therefore, even if specific colour pattern features might be under purifying selection among certain species (e.g., as a result of mimicry), this was not captured by latent variables capturing overarching differences between individuals and species in the data set. Our results indicate that aposematic species’ overall colour pattern phenotype might indeed be selected for less variability when compared to that of undefended species. However, our methodology does not address the possibility that specific colour pattern elements and signalling properties among aposematic species and putative mimics could be under purifying selection. Examples of this have been documented both within and between species of nudibranchs [3,47] and could apply to our dataset with representatives of a putative yellow-rim mimicry ring [96] (Fig. 1). This consideration is of broad relevance across all studies using methodology describing the cumulative colour pattern appearance of an animal, rather than specific colour pattern elements or body areas.

## Data accessibility

The data can be accessed on UQ’s e-space: https://doi.org/10.48610/a596710

## Authors’ contributions

CPvdB: Conceptualisation, data curation, formal analysis, funding acquisition, investigation, methodology, project administration, resources, software, validation, visualisation, writing – original draft, writing – review & editing. MS: Data analysis, writing - review and editing. JAE: project funding acquisition, writing - review and editing. BRD, CS, NW: investigation. KLC: Funding acquisition, project administration, resources, validation, writing - review & editing.

## Supporting information

electronic supplement

## Acknowledgements

We would like to thank various volunteers for assistance with image analysis, the High-Performance Computing (HPC) infrastructure at UQ (Wiener & Awoonga systems) and the infrastructure provided by Simone Blomberg, which contributed to the computing of image statistics.

