## Supplementary material for "Chemical defences indicate distinct colour patterns with reduced variability in aposematic nudibranchs": electronic supplement

### Table S1

Summary table of all individuals in the dataset used for this study.

| Species | Unpalatability (1-ED$_{50}$) | Toxicity (1-LD$_{50}$) | Picture taken | | Collection location | | | | | |
|---|---|---|---|---|---|---|---|---|---|---|
| | | | Animal in Laboratory | Animal in situ | Mackay, QLD | Sunshine Coast, QLD | Gold Coast, QLD | Nelson Bay, NSW | Cairns marine, QLD | Grand Total |
| *Aphelodoris varia* | 0 | 0 | 7 | 24 | 0 | 0 | 0 | 31 | 0 | 31 |
| *Chromodoris elisabethina* | 0.55 | 1 | 10 | 21 | 0 | 31 | 0 | 0 | 0 | 31 |
| *Chromodoris kuiteri* | 0.74 | 1 | 32 | 17 | 13 | 36 | 0 | 0 | 0 | 49 |
| *Chromodoris lochi* | 0.61 | 1 | 6 | 2 | 0 | 8 | 0 | 0 | 0 | 8 |
| *Chromodoris magnifica* | 0.65 | 1 | 11 | 3 | 8 | 4 | 0 | 0 | 2 | 14 |
| *Dendrodoris krusensterni* | 0 | 0 | 0 | 7 | 0 | 0 | 2 | 5 | 0 | 7 |
| *Discodoris sp.* | 0 | 0 | 3 | 7 | 0 | 1 | 0 | 9 | 0 | 10 |
| *Doriprismatica atromarginata* | 0.74 | 0.93 | 8 | 27 | 4 | 16 | 0 | 15 | 0 | 35 |
| *Glossodoris vespa* | 0.87 | 0.29 | 17 | 15 | 0 | 32 | 0 | 0 | 0 | 32 |
| *Hypselodoris bennetti* | 0.73 | 0.88 | 3 | 10 | 0 | 0 | 0 | 13 | 0 | 13 |
| *Phyllidia ocellata* | 0.63 | 0.4 | 9 | 23 | 4 | 28 | 0 | 0 | 0 | 32 |
| *Phyllidia varicosa* | 0.92 | 0.53 | 1 | 8 | 1 | 8 | 0 | 0 | 0 | 9 |
| *Phyllidiella pustulosa* | 0.86 | 0.14 | 22 | 18 | 7 | 33 | 0 | 0 | 0 | 40 |
| | | | 129 | 182 | | | | | | 311 |

**Figure S1**

Phylogenetic tree used in this study, modified from Cheney et al. 2014 as described in 'Methods'.

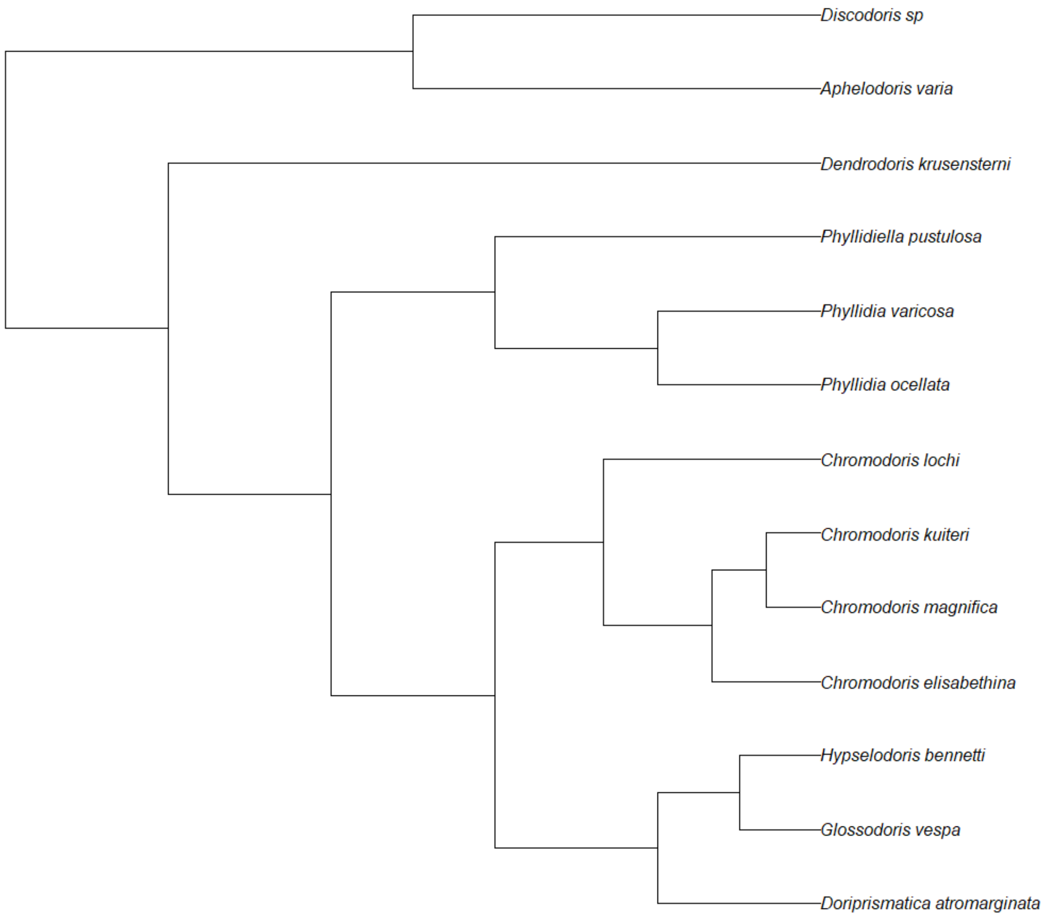

**Table S2**

Summary table of all >.6 Pearson correlation values between parameters with >0.4 loadings for each factor.

| FACTOR 1 | | | FACTOR 2 | | | FACTOR 3 | | |
|---|---|---|---|---|---|---|---|---|
| **Factor loading >0.4** | **Correlated Parameters (>.6)** | **Cor** | **Factor loading >0.4** | **Correlated Parameters (>.6)** | **Cor** | **Factor loading >0.4** | **Correlated Parameters (>.6)** | **Cor** |
| BSA.BMSL.Vrt | BSA.BML | 0.97 | CAA.Asp | Lum.mean.hrz | -0.67 | CAA.PT | CAA.PT.Hrz | 0.91 |
| | BSA.BMSL | 0.96 | | Lum.sd.hrz | -0.68 | | CAA.PT.Vrt | 0.91 |
| | BSA.BML.Hrz | 0.94 | CAA.Qt | CAA.Ht | 0.64 | CAA.Qc.Vrt | CAA.Jc | 0.73 |
| | BSA.BML.Vrt | 0.99 | | CAA.Qcpl | 0.83 | | CAA.Qc | 0.68 |
| | BSA.BMSL.Hrz | 0.92 | | CAA.Ht.Hrz | 0.65 | | CAA.Jc.Hrz | 0.72 |
| VCA.CVL.Hrz | Lum.sd | 0.67 | | CAA.Ht.Vrt | 0.62 | | CAA.Jc.Vrt | 0.73 |
| | Lum.sd.hrz | 0.66 | | CAA.Qt.Hrz | 0.88 | | CAA.Qc.Hrz | 0.97 |
| | VCA.CVL | 1.00 | | CAA.Qt.Vrt | 0.86 | Col.kurtosis.hrz | Col.CoV | 0.64 |
| | VCA.CVDmax | 0.67 | | CAA.Qcpl.Hrz | 0.83 | | Col.skew | 0.87 |
| | VCA.CVSsat | 0.68 | | CAA.Qcpl.Vrt | 0.83 | | Col.kurtosis | 0.98 |
| | VCA.CVL.Vrt | 1.00 | Lum.mean.vrt | Lum.mean | 0.83 | | Col.CoV.hrz | 0.62 |
| | VCA.CVDmax.Hrz | 0.68 | | Lum.sd | 0.67 | | Col.skew.hrz | 0.92 |
| | VCA.CVDmax.Vrt | 0.68 | | Lum.mean.hrz | 0.66 | | Col.skew.vrt | 0.75 |
| | VCA.CVSsat.Hrz | 0.69 | | Lum.sd.vrt | 0.84 | | Col.kurtosis.vrt | 0.85 |
| | VCA.CVSsat.Vrt | 0.69 | | CAA.C | 0.77 | Lum.kurtosis.hrz | Lum.skew | 0.83 |
| | BSA.BsL | 0.66 | | CAA.C.Hrz | 0.66 | | Lum.kurtosis | 0.95 |
| | BSA.BsSL | 0.70 | | CAA.C.Vrt | 0.80 | | Lum.skew.hrz | 0.91 |
| | BSA.BsL.Hrz | 0.67 | VCA.CVL.Hrz | Lum.sd | 0.67 | | Lum.kurtosis.vrt | 0.65 |
| | BSA.BsL.Vrt | 0.63 | | Lum.sd.hrz | 0.66 | Lum.mean.vrt | Lum.mean | 0.83 |
| | BSA.BsSL.Hrz | 0.70 | | VCA.CVL | 1.00 | | Lum.sd | 0.67 |
| | BSA.BsSL.Vrt | 0.67 | | VCA.CVDmax | 0.67 | | Lum.mean.hrz | 0.66 |

| | | |
|---|---|---|
| VCA.CVSL.Hrz | Col.mean | 0.61 |
| | Col.mean.vrt | 0.60 |
| | CAA.Hc | 0.62 |
| | CAA.Hc.Hrz | 0.62 |
| | CAA.Hc.Vrt | 0.61 |
| | VCA.CVSL | 1.00 |
| | VCA.CVSL.Vrt | 1.00 |
| | BSA.BCVL | 0.68 |
| | BSA.BCVSL | 0.65 |
| | BSA.BCVL.Hrz | 0.68 |
| | BSA.BCVL.Vrt | 0.67 |
| | BSA.BCVSL.Hrz | 0.65 |
| | BSA.BCVSL.Vrt | 0.65 |
| VCA.MDmax.Vrt | Col.mean | 0.78 |
| | Col.sd | 0.67 |
| | Col.mean.hrz | 0.75 |
| | Col.sd.hrz | 0.64 |
| | Col.mean.vrt | 0.80 |
| | Col.sd.vrt | 0.71 |
| | CAA.St.Hrz | 0.60 |
| | VCA.MDmax | 1.00 |
| | VCA.sDmax | 0.63 |
| | VCA.MSsat | 0.99 |
| | VCA.sSsat | 0.65 |
| | VCA.MDmax.Hrz | 1.00 |
| | VCA.sDmax.Hrz | 0.64 |
| | VCA.sDmax.Vrt | 0.64 |
| | VCA.MSsat.Hrz | 0.99 |
| | VCA.sSsat.Hrz | 0.65 |
| | VCA.MSsat.Vrt | 0.99 |
| | VCA.CVSsat | 0.68 |
| | VCA.CVL.Vrt | 1.00 |
| | VCA.CVDmax.Hrz | 0.68 |
| | VCA.CVDmax.Vrt | 0.68 |
| | VCA.CVSsat.Hrz | 0.69 |
| | VCA.CVSsat.Vrt | 0.69 |
| | BSA.BsL | 0.66 |
| | BSA.BsSL | 0.70 |
| | BSA.BsL.Hrz | 0.67 |
| | BSA.BsL.Vrt | 0.63 |
| | BSA.BsSL.Hrz | 0.70 |
| | BSA.BsSL.Vrt | 0.67 |
| VCA.CVS.Hrz | VCA.CVDmax | 0.62 |
| | VCA.CVSsat | 0.64 |
| | VCA.sS | 0.61 |
| | VCA.CVS | 1.00 |
| | VCA.CVDmax.Hrz | 0.62 |
| | VCA.CVDmax.Vrt | 0.62 |
| | VCA.CVSsat.Hrz | 0.64 |
| | VCA.CVSsat.Vrt | 0.64 |
| | VCA.sS.Hrz | 0.61 |
| | VCA.sS.Vrt | 0.61 |
| | VCA.CVS.Vrt | 1.00 |
| | BSA.BCVS | 0.78 |
| | BSA.BCVS.Hrz | 0.77 |
| | BSA.BCVS.Vrt | 0.78 |
| VCA.CVSL.Hrz | Col.mean | 0.61 |
| | Col.mean.vrt | 0.60 |
| | CAA.Hc | 0.62 |
| | CAA.Hc.Hrz | 0.62 |
| | Lum.sd.vrt | 0.84 |
| | CAA.C | 0.77 |
| | CAA.C.Hrz | 0.66 |
| | CAA.C.Vrt | 0.80 |

| | | | |
|---|---|---|---|
| VCA.sSsat.Vrt | 0.65 | CAA.Hc.Vrt | 0.61 |
| BSA.BCVL | 0.61 | VCA.CVSL | 1.00 |
| BSA.BMS | 0.65 | VCA.CVSL.Vrt | 1.00 |
| BSA.BCVL.Hrz | 0.61 | BSA.BCVL | 0.68 |
| BSA.BCVL.Vrt | 0.60 | BSA.BCVSL | 0.65 |
| BSA.BMS.Hrz | 0.64 | BSA.BCVL.Hrz | 0.68 |
| BSA.BMS.Vrt | 0.66 | BSA.BCVL.Vrt | 0.67 |
| | | BSA.BCVSL.Hrz | 0.65 |
| | | BSA.BCVSL.Vrt | 0.65 |

**Table S3**

Pairwise contrasts for the model investigating latent factor 1 expressed as the median differences between groups with different strengths of chemical defences (see 'Methods' for details). The effect size of pairwise differences increases with increasing deviation of such differences from zero, and the robustness of the result increases with decreasing degree of overlap of the 95% Credible Intervals (CIs) with zero.

| Factor 1: Contrasts | Difference | Lower CI | Upper CI |
|---|---|---|---|
| Undefended at 2cm - Toxic and moderately unpalatable at 2cm | 0.206 | -1.093 | 1.556 |
| Undefended at 2cm - Toxic and highly unpalatable at 2cm | -0.991 | -2.314 | 0.311 |
| Undefended at 2cm - Undefended at 10cm | -0.044 | -0.498 | 0.414 |
| Undefended at 2cm - Toxic and moderately unpalatable at 10cm | 0.355 | -1.024 | 1.676 |
| Undefended at 2cm - Toxic and highly unpalatable at 10cm | -0.644 | -1.981 | 0.733 |
| Toxic and moderately unpalatable at 2cm - Toxic and highly unpalatable at 2cm | -1.229 | -1.736 | -0.699 |
| Toxic and moderately unpalatable at 2cm - Undefended at 10cm | -0.257 | -1.700 | 1.073 |
| Toxic and moderately unpalatable at 2cm - Toxic and moderately unpalatable at 10cm | 0.145 | -0.222 | 0.448 |
| Toxic and moderately unpalatable at 2cm - Toxic and highly unpalatable at 10cm | -0.874 | -1.449 | -0.240 |
| Toxic and highly unpalatable at 2cm - Undefended at 10cm | 0.948 | -0.399 | 2.381 |
| Toxic and highly unpalatable at 2cm - Toxic and moderately unpalatable at 10cm | 1.363 | 0.735 | 1.913 |
| Toxic and highly unpalatable at 2cm - Toxic and highly unpalatable at 10cm | 0.346 | 0.043 | 0.703 |
| Undefended at 10cm - Toxic and moderately unpalatable at 10cm | 0.403 | -1.056 | 1.789 |
| Undefended at 10cm - Toxic and highly unpalatable at 10cm | -0.597 | -2.005 | 0.807 |
| Toxic and moderately unpalatable at 10cm - Toxic and highly unpalatable at 10cm | -1.014 | -1.665 | -0.326 |

**Table S4**

Pairwise variance contrasts for the model investigating latent factor 1 expressed as the median differences of the residual standard deviation on the original scale (back-transformed from the log scale) between groups with different strengths of chemical defences (see ‘Methods’ for details). The effect size of pairwise differences increases with increasing deviation of such differences from zero, and the robustness of the result increases with decreasing degree of overlap of the 95% Credible Intervals (CIs) with zero.

| Factor 1: Variance Contrasts | Difference | Lower CI | Upper CI |
|---|---|---|---|
| Undefended at 2cm - Toxic and moderately unpalatable at 2cm | 0.065 | -0.078 | 0.229 |
| Undefended at 2cm - Toxic and highly unpalatable at 2cm | -0.047 | -0.191 | 0.122 |
| Undefended at 2cm - Undefended at 10cm | -0.009 | -0.212 | 0.178 |
| Undefended at 2cm - Toxic and moderately unpalatable at 10cm | -0.027 | -0.184 | 0.096 |
| Undefended at 2cm - Toxic and highly unpalatable at 10cm | -0.035 | -0.157 | 0.068 |
| Toxic and moderately unpalatable at 2cm - Toxic and highly unpalatable at 2cm | -0.112 | -0.226 | 0.007 |
| Toxic and moderately unpalatable at 2cm - Undefended at 10cm | -0.074 | -0.270 | 0.089 |
| Toxic and moderately unpalatable at 2cm - Toxic and moderately unpalatable at 10cm | -0.092 | -0.354 | 0.125 |
| Toxic and moderately unpalatable at 2cm - Toxic and highly unpalatable at 10cm | -0.100 | -0.315 | 0.078 |
| Toxic and highly unpalatable at 2cm - Undefended at 10cm | 0.038 | -0.157 | 0.198 |
| Toxic and highly unpalatable at 2cm - Toxic and moderately unpalatable at 10cm | 0.020 | -0.228 | 0.209 |
| Toxic and highly unpalatable at 2cm - Toxic and highly unpalatable at 10cm | 0.012 | -0.230 | 0.214 |
| Undefended at 10cm - Toxic and moderately unpalatable at 10cm | -0.018 | -0.272 | 0.221 |
| Undefended at 10cm - Toxic and highly unpalatable at 10cm | -0.026 | -0.259 | 0.205 |
| Toxic and moderately unpalatable at 10cm - Toxic and highly unpalatable at 10cm | -0.008 | -0.183 | 0.176 |

**Table S5**

Pairwise contrasts for the model investigating latent factor 2 expressed as the median differences between groups with different strengths of chemical defences (see 'Methods' for details). The effect size of pairwise differences increases with increasing deviation of such differences from zero, and the robustness of the result increases with decreasing degree of overlap of the 95% Credible Intervals (CIs) with zero.

| Factor 2: Contrasts | Difference | Lower CI | Upper CI |
|---|---|---|---|
| Undefended at 2cm - Toxic and moderately unpalatable at 2cm | -1.021 | -2.595 | 0.637 |
| Undefended at 2cm - Toxic and highly unpalatable at 2cm | -0.960 | -2.478 | 0.717 |
| Undefended at 2cm - Undefended at 10cm | 0.064 | -0.269 | 0.393 |
| Undefended at 2cm - Toxic and moderately unpalatable at 10cm | -0.806 | -2.439 | 0.804 |
| Undefended at 2cm - Toxic and highly unpalatable at 10cm | -0.729 | -2.320 | 0.900 |
| Toxic and moderately unpalatable at 2cm - Toxic and highly unpalatable at 2cm | 0.031 | -0.824 | 0.933 |
| Toxic and moderately unpalatable at 2cm - Undefended at 10cm | 1.081 | -0.632 | 2.622 |
| Toxic and moderately unpalatable at 2cm - Toxic and moderately unpalatable at 10cm | 0.212 | 0.079 | 0.351 |
| Toxic and moderately unpalatable at 2cm - Toxic and highly unpalatable at 10cm | 0.263 | -0.571 | 1.228 |
| Toxic and highly unpalatable at 2cm - Undefended at 10cm | 1.029 | -0.667 | 2.549 |
| Toxic and highly unpalatable at 2cm - Toxic and moderately unpalatable at 10cm | 0.183 | -0.773 | 1.023 |
| Toxic and highly unpalatable at 2cm - Toxic and highly unpalatable at 10cm | 0.233 | 0.094 | 0.362 |
| Undefended at 10cm - Toxic and moderately unpalatable at 10cm | -0.864 | -2.474 | 0.789 |
| Undefended at 10cm - Toxic and highly unpalatable at 10cm | -0.800 | -2.299 | 0.952 |
| Toxic and moderately unpalatable at 10cm - Toxic and highly unpalatable at 10cm | 0.047 | -0.821 | 1.020 |

**Table S6**

Pairwise variance contrasts for the model investigating latent factor 2 expressed as the median differences of the residual standard deviation on the original scale (back-transformed from the log scale) between groups with different strengths of chemical defences (see 'Methods' for details). The effect size of pairwise differences increases with increasing deviation of such differences from zero, and the robustness of the result increases with decreasing degree of overlap of the 95% Credible Intervals (CIs) with zero.

| Factor 2: Variance Contrasts | Difference | Lower CI | Upper CI |
|---|---|---|---|
| Undefended at 2cm - Toxic and moderately unpalatable at 2cm | 0.397 | 0.141 | 0.743 |
| Undefended at 2cm - Toxic and highly unpalatable at 2cm | 0.314 | 0.056 | 0.656 |
| Undefended at 2cm - Undefended at 10cm | 0.047 | -0.270 | 0.337 |
| Undefended at 2cm - Toxic and moderately unpalatable at 10cm | -0.018 | -0.267 | 0.186 |
| Undefended at 2cm - Toxic and highly unpalatable at 10cm | -0.034 | -0.228 | 0.127 |
| Toxic and moderately unpalatable at 2cm - Toxic and highly unpalatable at 2cm | -0.083 | -0.211 | 0.052 |
| Toxic and moderately unpalatable at 2cm - Undefended at 10cm | -0.350 | -0.689 | -0.101 |
| Toxic and moderately unpalatable at 2cm - Toxic and moderately unpalatable at 10cm | -0.415 | -0.871 | -0.079 |
| Toxic and moderately unpalatable at 2cm - Toxic and highly unpalatable at 10cm | -0.431 | -0.850 | -0.134 |
| Toxic and highly unpalatable at 2cm - Undefended at 10cm | -0.267 | -0.611 | -0.015 |
| Toxic and highly unpalatable at 2cm - Toxic and moderately unpalatable at 10cm | -0.332 | -0.773 | -0.013 |
| Toxic and highly unpalatable at 2cm - Toxic and highly unpalatable at 10cm | -0.348 | -0.772 | -0.026 |
| Undefended at 10cm - Toxic and moderately unpalatable at 10cm | -0.064 | -0.457 | 0.299 |
| Undefended at 10cm - Toxic and highly unpalatable at 10cm | -0.081 | -0.447 | 0.264 |
| Toxic and moderately unpalatable at 10cm - Toxic and highly unpalatable at 10cm | -0.016 | -0.304 | 0.281 |

**Table S7**

Pairwise contrasts for the model investigating latent factor 3 expressed as the median differences between groups with different strength of chemical defences (see ‘Methods’ for details). Effect size of pairwise differences increases with increasing deviation of such differences from zero, and the robustness of the result increases with decreasing degree of overlap of the 95% Credible Intervals (CIs) with zero.

| Factor 3: Contrasts | Difference | Lower CI | Upper CI |
|---|---|---|---|
| Undefended at 2cm - Toxic and moderately unpalatable at 2cm | 0.052 | -1.163 | 1.143 |
| Undefended at 2cm - Toxic and highly unpalatable at 2cm | -0.104 | -1.288 | 1.030 |
| Undefended at 2cm - Undefended at 10cm | 0.695 | 0.280 | 1.124 |
| Undefended at 2cm - Toxic and moderately unpalatable at 10cm | 0.666 | -0.476 | 1.793 |
| Undefended at 2cm - Toxic and highly unpalatable at 10cm | 0.587 | -0.645 | 1.640 |
| Toxic and moderately unpalatable at 2cm - Toxic and highly unpalatable at 2cm | -0.163 | -0.765 | 0.456 |
| Toxic and moderately unpalatable at 2cm - Undefended at 10cm | 0.633 | -0.433 | 1.736 |
| Toxic and moderately unpalatable at 2cm - Toxic and moderately unpalatable at 10cm | 0.610 | 0.352 | 0.862 |
| Toxic and moderately unpalatable at 2cm - Toxic and highly unpalatable at 10cm | 0.528 | -0.016 | 1.100 |
| Toxic and highly unpalatable at 2cm - Undefended at 10cm | 0.805 | -0.308 | 1.903 |
| Toxic and highly unpalatable at 2cm - Toxic and moderately unpalatable at 10cm | 0.773 | 0.189 | 1.299 |
| Toxic and highly unpalatable at 2cm - Toxic and highly unpalatable at 10cm | 0.692 | 0.443 | 0.936 |
| Undefended at 10cm - Toxic and moderately unpalatable at 10cm | -0.020 | -1.174 | 0.976 |
| Undefended at 10cm - Toxic and highly unpalatable at 10cm | -0.110 | -1.203 | 0.961 |
| Toxic and moderately unpalatable at 10cm - Toxic and highly unpalatable at 10cm | -0.083 | -0.567 | 0.422 |

**Table S8**

Pairwise variance contrasts for the model investigating latent factor 3 expressed as the median differences of the residual standard deviation on the original scale (back-transformed from the log scale) between groups with different strengths of chemical defences (see ‘Methods’ for details). The effect size of pairwise differences increases with increasing deviation of such differences from zero, and the robustness of the result increases with decreasing degree of overlap of the 95% Credible Intervals (CIs) with zero.

| **Factor 3: Variance Contrasts** | **Difference** | **Lower CI** | **Upper CI** |
|---|---|---|---|
| Undefended at 2cm - Toxic and moderately unpalatable at 2cm | 0.338 | -0.059 | 0.945 |
| Undefended at 2cm - Toxic and highly unpalatable at 2cm | 0.287 | -0.099 | 0.893 |
| Undefended at 2cm - Undefended at 10cm | 0.218 | -0.088 | 0.670 |
| Undefended at 2cm - Toxic and moderately unpalatable at 10cm | 0.187 | -0.036 | 0.473 |
| Undefended at 2cm - Toxic and highly unpalatable at 10cm | 0.188 | -0.003 | 0.447 |
| Toxic and moderately unpalatable at 2cm - Toxic and highly unpalatable at 2cm | -0.052 | -0.295 | 0.194 |
| Toxic and moderately unpalatable at 2cm - Undefended at 10cm | -0.120 | -0.484 | 0.164 |
| Toxic and moderately unpalatable at 2cm - Toxic and moderately unpalatable at 10cm | -0.151 | -0.694 | 0.227 |
| Toxic and moderately unpalatable at 2cm - Toxic and highly unpalatable at 10cm | -0.150 | -0.641 | 0.181 |
| Toxic and highly unpalatable at 2cm - Undefended at 10cm | -0.069 | -0.435 | 0.225 |
| Toxic and highly unpalatable at 2cm - Toxic and moderately unpalatable at 10cm | -0.100 | -0.617 | 0.242 |
| Toxic and highly unpalatable at 2cm - Toxic and highly unpalatable at 10cm | -0.099 | -0.618 | 0.257 |
| Undefended at 10cm - Toxic and moderately unpalatable at 10cm | -0.031 | -0.434 | 0.282 |
| Undefended at 10cm - Toxic and highly unpalatable at 10cm | -0.030 | -0.398 | 0.260 |
| Toxic and moderately unpalatable at 10cm - Toxic and highly unpalatable at 10cm | 0.001 | -0.277 | 0.284 |

**Table S9**

Coefficient estimates of the model investigating the scores for latent factor 1 between species of nudibranchs with different levels of chemical defences (N = 13, R^2 = 0.58). Estimates are based on a Student distribution with an identity link for the mean of the response distribution and a log link for its residual standard deviation (Sigma). The estimate is more likely to be non-zero when the credible intervals do not overlap with zero.

| Coefficient | Mean | M. Error | 95% CIs | |
|---|---|---|---|---|
| | | | Low | High |
| **Group-level effects** | | | | |
| *Phylogenesis* | | | | |
| Sd [Intercept] | 0.63 | 0.28 | 0.09 | 1.23 |
| *Species of nudibranch* | | | | |
| Sd [Intercept: Distance 2 cm] | 0.24 | 0.18 | 0.01 | 0.67 |
| Sd [Distance 10 cm] | 0.31 | 0.11 | 0.14 | 0.57 |
| Sigma: Sd [Intercept: Distance 2 cm] | 0.07 | 0.05 | 0.00 | 0.2 |
| Sigma: Sd [Distance 10 cm] | 0.08 | 0.07 | 0.00 | 0.26 |
| **Population-level effects** | | | | |
| Intercept [Undefended, 2 cm] | -0.39 | 0.49 | -1.40 | 0.58 |
| Sigma_Intercept [Undefended, 2 cm] | -0.72 | 0.14 | -0.98 | -0.45 |
| Chemical defence [Toxic and moderately unpalatable] | -0.22 | 0.66 | -1.62 | 1.05 |
| Chemical defence [Toxic and highly unpalatable] | 0.99 | 0.65 | -0.35 | 2.3 |
| Distance [10 cm] | 0.05 | 0.24 | -0.41 | 0.51 |
| Chemical defence [Toxic and moderately unpalatable] x Distance [10 cm] | -0.19 | 0.29 | -0.75 | 0.39 |
| Chemical defence [Toxic and highly unpalatable] x Distance [10 cm] | -0.40 | 0.28 | -0.96 | 0.16 |
| Sigma: Chemical defence [Toxic and moderately unpalatable] | -0.14 | 0.16 | -0.46 | 0.18 |
| Sigma: Chemical defence [Toxic and highly unpalatable] | 0.10 | 0.16 | -0.22 | 0.39 |
| Sigma: Distance [10cm] | 0.01 | 0.20 | -0.37 | 0.4 |
| Sigma: Chemical defence [Toxic and moderately unpalatable] x Distance [10 cm] | 0.03 | 0.24 | -0.44 | 0.5 |
| Sigma: Chemical defence [Toxic and highly unpalatable] x Distance [10 cm] | 0.05 | 0.22 | -0.39 | 0.49 |

**Table S10**

Coefficient estimates of the model investigating the scores for latent factor 2 between species of nudibranchs with different levels of chemical defences (N = 13, R^2 = 0.66). Estimates are based on a Student distribution with an identity link for the mean of the response distribution and a log link for its residual standard deviation (Sigma). The estimate is more likely to be non-zero when the credible intervals do not overlap with zero.

| Coefficient | Mean | M. Error | 95% CIs | |
|---|---|---|---|---|
| | | | Low | High |
| **Group-level effects** | | | | |
| *Phylogenesis* | | | | |
| Sd [Intercept] | 0.58 | 0.44 | 0.02 | 1.62 |
| *Species of nudibranch* | | | | |
| Sd [Intercept: Distance 2 cm] | 0.59 | 0.23 | 0.14 | 1.07 |
| Sd [Distance 10 cm] | 0.08 | 0.06 | 0.00 | 0.22 |
| Sigma: Sd [Intercept: Distance 2 cm] | 0.21 | 0.09 | 0.07 | 0.42 |
| Sigma: Sd [Distance 10 cm] | 0.12 | 0.10 | 0.00 | 0.37 |
| **Population-level effects** | | | | |
| Intercept [Undefended, 2 cm] | -0.78 | 0.60 | -1.97 | 0.45 |
| Sigma_Intercept [Undefended, 2 cm] | -0.37 | 0.21 | -0.78 | 0.05 |
| Chemical defence [Toxic and moderately unpalatable] | 0.99 | 0.82 | -0.74 | 2.54 |
| Chemical defence [Toxic and highly unpalatable] | 0.95 | 0.81 | -0.69 | 2.5 |
| Distance [10 cm] | -0.06 | 0.17 | -0.40 | 0.26 |
| Chemical defence [Toxic and moderately unpalatable] x Distance [10 cm] | -0.15 | 0.18 | -0.50 | 0.21 |
| Chemical defence [Toxic and highly unpalatable] x Distance [10 cm] | -0.17 | 0.18 | -0.51 | 0.2 |
| Sigma: Chemical defence [Toxic and moderately unpalatable] | -0.82 | 0.24 | -1.30 | -0.3 |
| Sigma: Chemical defence [Toxic and highly unpalatable] | -0.58 | 0.23 | -1.05 | -0.1 |
| Sigma: Distance [10cm] | -0.07 | 0.22 | -0.49 | 0.38 |
| Sigma: Chemical defence [Toxic and moderately unpalatable] x Distance [10 cm] | 0.08 | 0.27 | -0.46 | 0.59 |
| Sigma: Chemical defence [Toxic and highly unpalatable] x Distance [10 cm] | 0.11 | 0.25 | -0.39 | 0.58 |

**Table S11**

Coefficient estimates of the model investigating the scores for latent factor 3 between species of nudibranchs with different levels of chemical defences (N = 13, R^2 = 0.37). Estimates are based on a Student distribution with an identity link for the mean of the response distribution and a log link for its residual standard deviation (Sigma). The estimate is more likely to be non-zero when the credible intervals do not overlap with zero.

| Coefficient | Mean | M. Error | 95% CIs | |
|---|---|---|---|---|
| | | | Low | High |
| **Group-level effects** | | | | |
| *Phylogenesis* | | | | |
| Sd [Intercept] | 0.43 | 0.29 | 0.02 | 1.06 |
| *Species of nudibranch* | | | | |
| Sd [Intercept: Distance 2 cm] | 0.38 | 0.21 | 0.02 | 0.77 |
| Sd [Distance 10 cm] | 0.24 | 0.08 | 0.13 | 0.43 |
| Sigma: Sd [Intercept: Distance 2 cm] | 0.50 | 0.14 | 0.28 | 0.85 |
| Sigma: Sd [Distance 10 cm] | 0.27 | 0.15 | 0.02 | 0.6 |
| **Population-level effects** | | | | |
| Intercept [Undefended, 2 cm] | 0.23 | 0.44 | -0.65 | 1.07 |
| Sigma_Intercept [Undefended, 2 cm] | -0.51 | 0.36 | -1.19 | 0.21 |
| Chemical defence [Toxic and moderately unpalatable] | -0.05 | 0.57 | -1.20 | 1.12 |
| Chemical defence [Toxic and highly unpalatable] | 0.12 | 0.57 | -1.03 | 1.29 |
| Distance [10 cm] | -0.70 | 0.21 | -1.12 | -0.28 |
| Chemical defence [Toxic and moderately unpalatable] x Distance [10 cm] | 0.08 | 0.25 | -0.41 | 0.56 |
| Chemical defence [Toxic and highly unpalatable] x Distance [10 cm] | 0.00 | 0.25 | -0.50 | 0.49 |
| Sigma: Chemical defence [Toxic and moderately unpalatable] | -0.72 | 0.44 | -1.60 | 0.16 |
| Sigma: Chemical defence [Toxic and highly unpalatable] | -0.56 | 0.42 | -1.40 | 0.25 |
| Sigma: Distance [10cm] | -0.41 | 0.29 | -1.00 | 0.18 |
| Sigma: Chemical defence [Toxic and moderately unpalatable] x Distance [10 cm] | 0.04 | 0.38 | -0.72 | 0.76 |
| Sigma: Chemical defence [Toxic and highly unpalatable] x Distance [10 cm] | 0.05 | 0.34 | -0.64 | 0.71 |
